# Biogenic Amine Neurotransmitters Promote Eicosanoid Production And Protein Homeostasis

**DOI:** 10.1101/2020.02.25.964189

**Authors:** Kishore K. Joshi, Tarmie L. Matlack, Stephanie Pyonteck, Ralph Menzel, Christopher Rongo

## Abstract

Multicellular organisms use multiple pathways to restore protein homeostasis (proteostasis) in response to adverse physiological conditions, changing environment, and developmental aging. The nervous system can regulate proteostasis in different tissues, but it is unclear how it mobilizes proteostasis pathways to offset physiological decline. Here we show that *C. elegans* employs the humoral biogenic amine neurotransmitters dopamine, serotonin, and tyramine to regulate proteostasis and the activity of the Ubiquitin Proteasome System (UPS) in epithelial tissues. Mutants for biogenic amine synthesis show decreased poly-ubiquitination and turnover of a GFP-based UPS substrate. Using RNA-seq, we determined the expression profile of genes regulated by biogenic amine signaling. We find that biogenic amines promote the expression of a subset of cytochrome P450 monooxygenases involved in eicosanoid production from polyunsaturated fatty acids (PUFAs). Mutants for these P450s share the same UPS phenotype observed in biogenic amine mutants. The production of n-3 PUFAs is required for UPS substrate turnover, whereas mutants that accumulate n-3 PUFAs show accelerated turnover of this GFP-based substrate. Our results suggest that neurosecretory sensory neurons release biogenic amines to modulate the lipid signaling profile, which in turn activates stress response pathways to maintain proteostasis.

## INTRODUCTION

Cells employ multiple signaling pathways and mechanisms for maintaining the folded state and function of their proteins, a process termed protein homeostasis (proteostasis) (Balch, Morimoto et al., 2008, Hipp, Kasturi et al., 2019). Challenges to proteostasis can come from changes in the environment (e.g., extreme temperature), altered internal physiology (e.g., reactive oxygen species or ROS from mitochondrial dysfunction), and physiologically-associated decline due to aging (Shore & Ruvkun, 2013, Warnatsch, Bergann et al., 2013). Proteostasis response pathways offset these challenges by either restoring proper protein folding or removing unfolded or damaged proteins from cells. The failure to maintain proteostasis in the face of such challenges results in physiological decline from the accumulation of oxidized and unfolded proteins.

One of the key pathways that removes damaged and unfolded proteins is the Ubiquitin Proteasome System (UPS), which covalently attaches chains of the small protein ubiquitin to lysine side chains of protein substrates (Ciechanover & Stanhill, 2014). E3-type ubiquitin ligases either specifically recognize protein substrates or recognize unfolded proteins with the help of chaperone adapters, attaching single ubiquitin protein moieties (mono-ubiquitination) to these substrates (Zheng & Shabek, 2017). E4-type ubiquitin ligases recognize mono-ubiquitinated proteins and attach additional ubiquitin molecules to the initial ubiquitin, resulting in poly-ubiquitin chains (Hoppe, 2005). Poly-ubiquitinated substrates are then degraded by proteolysis via the 26S proteasome (Bard, Goodall et al., 2018). Aggregation-prone poly-ubiquitinated proteins are sometimes resistant to proteasomal degradation and must be recognized and removed by additional proteostasis response pathways, including the autophagy pathway (Ciechanover & Kwon, 2015, Cortes & La Spada, 2015, Kirkin, McEwan et al., 2009, Lim & Yue, 2015, Ryno, Wiseman et al., 2013). It is unclear how these different pathways are coordinated or how some substrates are preferred by one pathway over another (Kocaturk & Gozuacik, 2018).

Multicellularity affords organisms the ability to employ different proteostasis pathways depending on the type of tissue or the particular proteostasis challenge. The nervous system coordinates physiology, including proteostasis, throughout multiple tissues through humoral signals released in the body (Balch et al., 2008, O’Brien & van Oosten-Hawle, 2016, van Oosten-Hawle & Morimoto, 2014). Biogenic amines – a family of neurotransmitters formed from the decarboxylation of amino acids – can act as such humoral signals when released into the body from neurosecretory cells or even non-neuronal cells (Morgese & Trabace, 2019, Ng, Papandreou et al., 2015, Zeng, Zhang et al., 2007). In the nematode *C. elegans*, thermosensory neurons release the biogenic amine neurotransmitter serotonin, which is then able to activate the expression of heat shock proteins (HSPs) in distal tissues (Prahlad, Cornelius et al., 2008, Tatum, Ooi et al., 2015). Similarly, octopaminergic sensory neurons modulate the Unfolded Protein Response (UPR) in distal tissues (Sun, Liu et al., 2012, Sun, Singh et al., 2011). It is unclear whether biogenic amines directly mediate their effects on target tissues or act through additional signaling intermediates to modulate proteostasis.

We previously identified novel regulators of proteostasis in *C. elegans* by using Ub^G76V^-GFP, a Ubiquitin Fusion Degradation (UFD) substrate expressed from a transgene, to monitor UPS activity (Joshi, Matlack et al., 2016, Liu, Rogers et al., 2011). Ub^G76V^-GFP contains a non-cleavable ubiquitin placed amino-terminal to GFP (Liu et al., 2011, Segref, Torres et al., 2011), thereby mimicking a mono-ubiquitinated protein. E3- and E4-type ubiquitin ligases in the UFD complex attach additional ubiquitins to the K48 residue of Ub^G76V^-GFP, resulting in poly-ubiquitination and proteasomal degradation of this reporter (Butt, Khan et al., 1988, Dantuma, Lindsten et al., 2000, Johnson, Ma et al., 1995). Ub^G76V^-GFP degradation was previously monitored in *C. elegans* intestinal or hypodermal epithelia when this UFD substrate was expressed from either the *sur-5* or *col-19* promoter, respectively (Liu et al., 2011, Segref et al., 2011). Using this reporter, we found that the biogenic amine dopamine non-autonomously promotes UPS activity in intestinal and hypodermal epithelia in *C. elegans* (Joshi et al., 2016). Whether *C. elegans* employs other biogenic amines to regulate the UPS is unknown.

*C. elegans* has four biogenic amine neurotransmitters: serotonin (5HT), dopamine (DA), octopamine (OA), and tyramine (TA) (Chase & Koelle, 2007). The *tph-1* gene encodes the *C. elegans* tryptophan hydroxylase, the key enzyme for 5HT biosynthesis from tryptophan (Sze, Victor et al., 2000). The *cat-2* gene encodes the *C. elegans* tyrosine hydroxylase, which catalyzes the rate-limiting step in DA synthesis from tyrosine (Lints & Emmons, 1999). The biogenic amines TA and OA are also synthesized from tyrosine but along a different biochemical pathway (Alkema, Hunter-Ensor et al., 2005). OA is made from TA, and mutants for the *tbh-1* tyramine beta hydroxylase fail to produce OA, whereas mutants for *tdc-1* tyrosine decarboxylase produce neither OA nor TA. Although TA and OA are synthesized along the same pathway, they have independent functions. All four neurotransmitters modulate multiple neural circuits involved in locomotion, egg-laying, and feeding behaviors in *C. elegans* (Chase & Koelle, 2007). All four are involved in how nematodes respond to food, although the full scope of their signaling is not known.

Here we used the Ub^G76V^-GFP reporter to examine UPS activity in mutants that fail to synthesize the different biogenic amines in *C. elegans*. We found that the biogenic amine neurotransmitters DA, 5HT, and TA modulate Ub^G76V^-GFP turnover in epithelia, as hypodermal epithelia in mutants lacking DA, 5HT, or TA show increased stability of the otherwise unstable Ub^G76V^-GFP reporter protein. Using RNA-seq, we determined the expression profile of biogenic amine mutants and found that biogenic amine signaling promotes the expression of a specific subclass of cytochrome P450 monooxygenases (P450) involved in the production of eicosanoids from poly-unsaturated fatty acids (PUFAs). Mutants for one of these regulated P450s, *cyp-34A4,* showed stabilized Ub^G76V^-GFP, similar to the phenotype observed in biogenic amine mutants. Consistent with our expression profile analysis, we found that biogenic amine mutants have depressed levels of eicosanoids. As eicosanoids are derived from n-3 or n-6 long chain PUFAs, we examined mutants impaired for PUFA synthesis and found that n-3 lipids are necessary and sufficient to promote Ub^G76V^-GFP turnover. Our results suggest that neurons use biogenic amine neurotransmitters as neurohormonal signals to modulate eicosanoid production from PUFAs, and that these eicosanoids in turn regulate proteostasis response pathways in non-neuronal tissues.

## RESULTS

### The Ub^G76V^-GFP transgene reports UPS activity

We sought to examine changes in the hypodermis – the epithelial layer surrounding the nematode body cavity and responsible for body growth and synthesis of the outer barrier of the collagenous cuticle. The *odIs77* integrated transgene expresses both the chimeric Ub^G76V^-GFP protein and mRFP in the hypodermis from the *col-19* promoter (Liu et al., 2011). The levels of Ub^G76V^-GFP, which is a UPS substrate, can be compared to the levels of mRFP, which remains stable (Fig. 1A). The mRFP allows researchers to quickly rule out changes in transgene expression versus changes in Ub^G76V^-GFP protein degradation. While mRFP protein remains stable, the Ub^G76V^-GFP protein is rapidly degraded as nematodes move from the first day of adulthood (Fig. 1B,D; L4+24h) into the second day of adulthood (Fig. 1C,E; L4+48h), and this turnover is largely prevented when either the UFD complex or the proteasome is inhibited (Joshi et al., 2016). The ratio of Ub^G76V^-GFP to mRFP levels inversely reflect UPS activity; thus, hypodermal UPS activity increases significantly as nematodes enter day two of adulthood, a period of peak fecundity.

**Figure 1.**
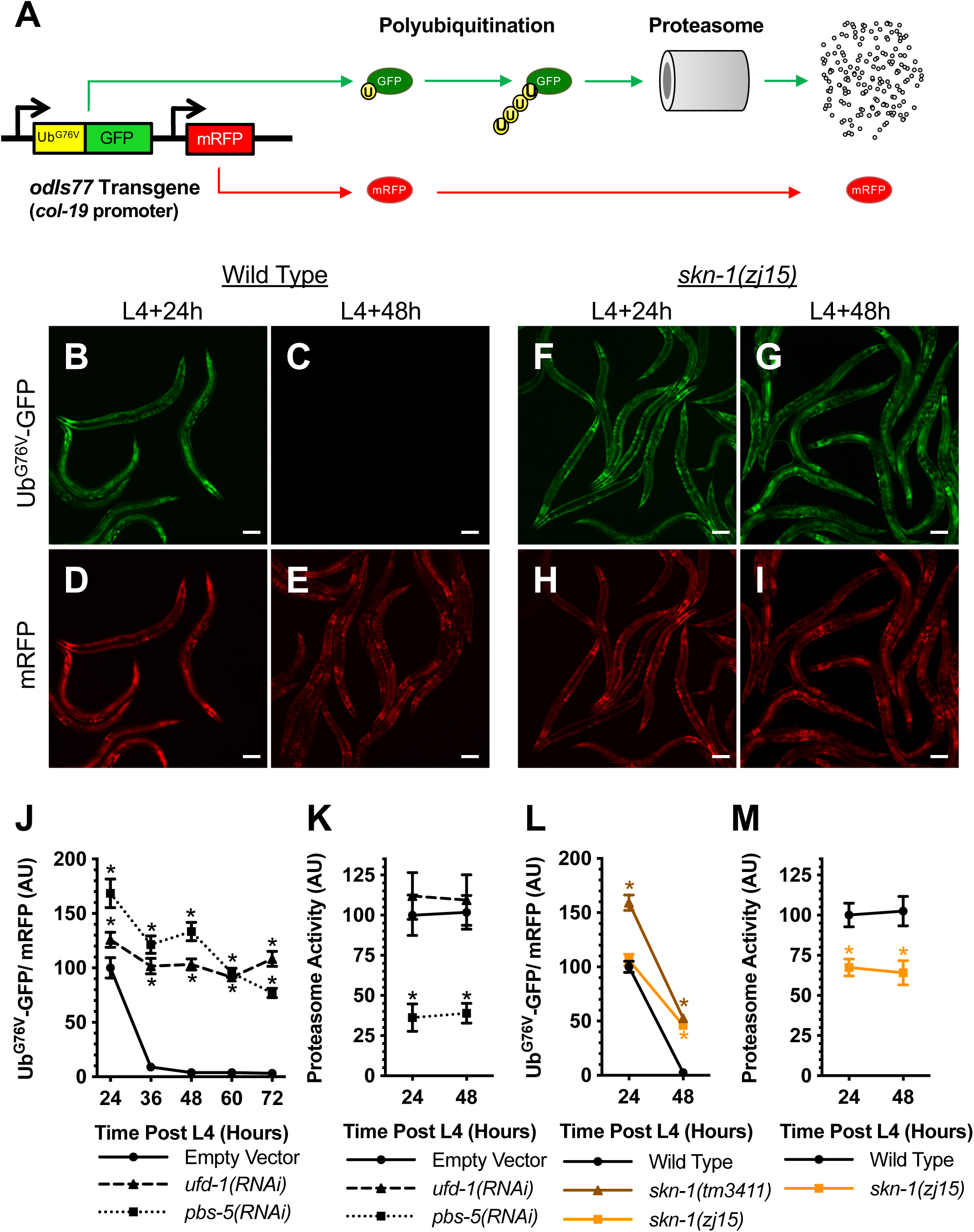
The Ub^G76V^-GFP transgene reports UPS activity. A Schematic representation of the Ub^G76V^-GFP chimeric reporter (yellow for ubiquitin and green for GFP) and its associated mRFP internal control (red). The Ub^G76V^-GFP chimera, which contains a mutation in the terminal residue of ubiquitin, cannot be cleaved. The resulting protein is a substrate for poly-ubiquitination (indicated by the circles labeled with “u”) and degradation by the proteasome. High levels of reporter poly-ubiquitination and/or proteasome activity result in little or no GFP fluorescence. B,C Expression via the *col-19* promoter of Ub^G76V^-GFP in *C. elegans* hypodermis from a single integrated transgene at (B) L4+24 hours and (C) L4+48 hours. Wild-type animals are shown. Bar, 100 microns. D,E Expression of mRFP in the same animals as (B,C) at (D) L4+24 hours and (E) L4+48 hours. F,G Expression via the *col-19* promoter of Ub^G76V^-GFP in *C. elegans* hypodermis from a single integrated transgene at (B) L4+24 hours and (C) L4+48 hours. Mutants for *skn-1(zj15)* are shown. Bar, 100 microns. H,I Expression of mRFP in the same animals as (F,G) at (H) L4+24 hours and (I) L4+48 hours. J Quantified fluorescence of Ub^G76V^-GFP normalized to mRFP in the hypodermis of animals from the indicated time point (in hours) after the L4 stage. Animals have been exposed to the indicated knockdown RNAi bacterial strains or a strain that only contains the empty RNAi vector since hatching. *P<0.001, ANOVA with Dunnetts posthoc comparison to the wild-type control equivalent time point. N=20 animals per genotype and time point. Error bars indicate SEM. K Quantified epoxomicin-sensitive proteasome activity (as measured through the turnover of fluorescent chymotrypsin substrate) from lysates of animals treated as in (J). *P<0.001, ANOVA with Dunnetts posthoc comparison to the empty RNAi vector control. N=3 trials. Error bars indicate SEM. L Quantified fluorescence of Ub^G76V^-GFP normalized to mRFP in the hypodermis of animals from the indicated time point (in hours) and the indicated genotypes after the L4 stage, as per (J). M Quantified epoxomicin-sensitive proteasome activity, as per (K).

We revalidated our transgenic reporter by examining transgenic animals impaired for UPS activity. Ub^G76V^-GFP is polyubiquitinated by the UFD complex and degraded by the 26S proteasome (Joshi et al., 2016, Liu et al., 2011, Segref et al., 2011). Eggs harboring the *odIs77* transgene were hatched on bacterial lawns expressing RNAi constructs for either the UFD complex subunit *ufd-1*, the proteasome complex core particle subunit *pbs-5*, or an empty vector as control. Ub^G76V^-GFP levels were elevated relative to control in the *ufd-1* and *pbs-5* RNAi knockdown experiments by both day 1 and day 2 adulthood (Fig. 1J), consistent with UPS-mediated turnover of this chimeric reporter protein. We directly examined proteasome activity by generating lysates from the animals and incubating them with the fluorescent chymotryptic substrate Suc-Leu-Leu-Val-Tyr-AMC in the presence or absence of the proteasome inhibitor epoxomicin. Knockdown of the proteasome subunit *pbs-5* resulted in diminished proteasome activity relative to empty vector control (Fig. 1K). Knockdown of *ufd-1*, a gene involved in polyubiquitination but not proteasome catalytic activity, did not diminish proteasome activity, as expected (Fig. 1K). These results confirm that the increased levels of the Ub^G76V^-GFP chimeric reporter reflect decreases in UFD activity (and thus polyubiquitination) or proteasome activity.

The transcription factor SKN-1, which is part of the oxidative stress response and homologous to human Nrf2, directly promotes the expression of proteasome subunit genes (An & Blackwell, 2003, Keith, Maddux et al., 2016, Li, Matilainen et al., 2011, Niu, Lu et al., 2011). We examined Ub^G76V^-GFP levels in *skn-1(zj15)*, a fertile hypomorphic allele, and *skn-1(tm3411)*, a null deletion allele that results in sterile animals (Ruf, Holzem et al., 2013, Tang, Dodd et al., 2015). Ub^G76V^-GFP was present in *skn-1(zj15)* in day 1 adults (Fig. 1F,H) and remained stable in day 2 (Fig. 1G,I) adults. Ub^G76V^-GFP was similarly stable in *skn-1(tm3411)* null mutants (Fig. 1L). As *skn-1(zj15)* animals are fertile and do not require a balancer chromosome, we were able to collect large numbers of synchronized animals and measure proteasome activity. Mutants for *skn-1(zj15)* showed about a 30% decrease in proteasome activity relative to wild type in both day 1 and day 2 adults (Fig. 1M), consistent with the decreased levels of proteasome subunit expression (Keith et al., 2016, Li et al., 2011, Niu et al., 2011) and stabilized UPS reporter (Fig. 1L) observed in mutants. In addition to providing a feedback loop to activate proteasome expression during proteotoxic stress, SKN-1 promotes baseline UPS activity.

### Biogenic amine signaling modulates protein turnover

In order to test each biogenic amine for a role in regulating proteostasis, we examined mutants for specific biosynthetic enzymes (Fig. 2A). We introduced the *odIs77* transgene into *cat-2(e1112)* mutants, which have a nonsense mutation in *cat-2* and fail to synthesize DA, *tph-1(mg280)* mutants, which harbor a deletion of *tph-1* sequences and fail to synthesize 5HT, *tdc-1(ok914)* mutants, which contain a deletion of *tdc-1* sequences and fail to synthesize TA and OA, and *tbh-1(ok1196)* mutants, which harbor a deletion of *tbh-1* sequences and fail to synthesize OA (Alkema et al., 2005, Lints & Emmons, 1999, Sulston, Dew et al., 1975, Sze et al., 2000). In addition to qualitatively assessing fluorescence photomicrographs (Fig. 2C-F), we quantified Ub^G76V^-GFP and mRFP levels for each genotype (Fig. 2B,K). Ub^G76V^-GFP levels in *tph-1* and *tdc-1* mutants were similar to those in wild type in day 1 adults but did not diminish to the same extent at subsequent time points (Fig. 2B), suggesting reduced turnover. Ub^G76V^-GFP levels in *cat-2* were about 50% higher than those in wild type in day 1 adults and remained higher throughout our time course analysis (Fig. 2B-F). We observed no difference between *tbh-1* mutants and wild type, suggesting that OA is not required to regulate Ub^G76V^-GFP turnover (Fig. 2K). Investigation of additional alleles of each gene at day 2 (L4+48h) were consistent and confirmed the above findings (Fig. 2K). To confirm that any observed regulation of Ub^G76V^-GFP occurs at the level of protein stability rather than transcription, we quantified Ub^G76V^-GFP and mRFP mRNA levels and detected no change between the different genotypes and developmental time points (data not shown), consistent with a post-transcriptional explanation for the observed differences in protein levels. Our results indicate that UPS activity is decreased in mutants that fail to synthesize the biogenic amines DA, 5HT, and TA, but not in mutants that fail to synthesize only OA.

**Figure 2.**
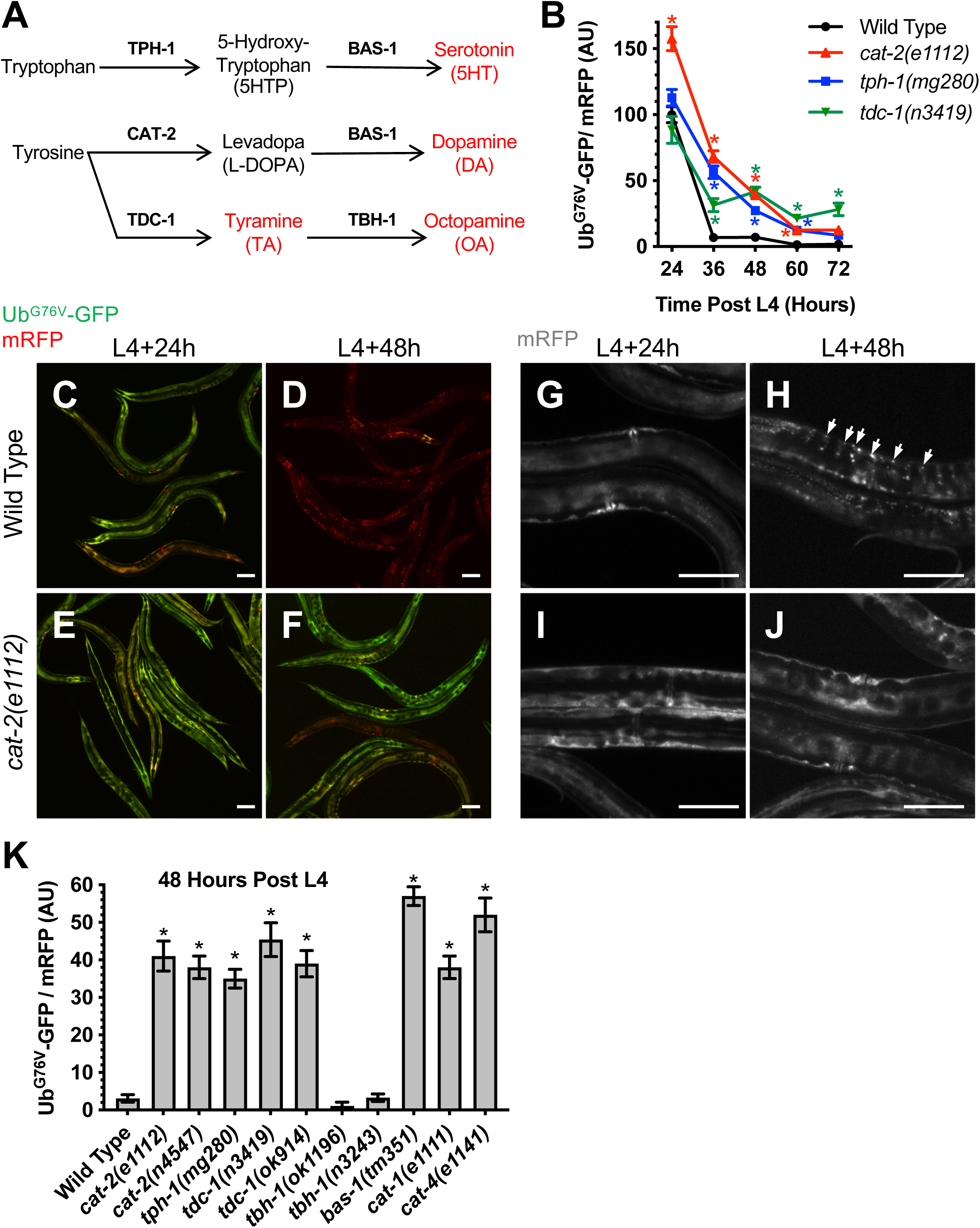
Biogenic amine signaling modulates protein turnover. A Schematic representation of the biogenic amine synthesis pathways in *C. elegans*. Final biogenic amine neurotransmitter products are shown in red. The gene encoding each biosynthesis enzyme in the pathway is indicated over the arrow representing its catalyzed reaction. B Quantified fluorescence of Ub^G76V^-GFP normalized to mRFP in the hypodermis of animals of the specified genotype from the indicated time point (in hours) after the L4 stage. *P<0.001, ANOVA with Dunnetts posthoc comparison to the wild-type control equivalent time point. N=20 animals per genotype and time point. Error bars indicate SEM. C,D Co-expression via the *col-19* promoter of Ub^G76V^-GFP (green) and mRFP (red) in *C. elegans* hypodermis from a single integrated transgene at (C) L4+24 hours and (D) L4+48 hours. Green and red channels are merged. Wild-type animals are shown. Bar, 100 microns. E,F Similar analysis as in (C,D) for *cat-2(e1112)* mutants. G,H Expression of mRFP via the *col-19* promoter in *C. elegans* hypodermis at (G) L4+24 hours or (H) L4+48 hours. Wild-type animals are shown. Puncta containing mRFP aggregates are indicated by arrows. Bar, 100 microns. I,J Similar analysis as in (G,H) for *cat-2(e1112)* mutants. K Quantified fluorescence of Ub^G76V^-GFP normalized to mRFP in the hypodermis of animals of the indicated genotype at L4+48 hours. *P<0.001, ANOVA with Dunnetts posthoc comparison to the wild-type control equivalent time point. N=20 animals per genotype and time point. Error bars indicate SEM.

We also examined mutants that impair signaling of more than one biogenic amine. BAS-1 encodes an aromatic amino acid decarboxylase required for the synthesis of both DA and 5HT, CAT-1 encodes a monoamine transporter required for loading synaptic vesicles with likely all biogenic amines, and CAT-4 encodes a GTP cyclohydrolase required for the synthesis of multiple small compounds including DA and 5HT (Duerr, Frisby et al., 1999, Hare & Loer, 2004, Loer & Kenyon, 1993, Sulston et al., 1975). Ub^G76V^-GFP levels in null mutants for all three of these genes were similar to those in wild type in day 1 adults but remained stable in day 2 adults (Fig. 2K), suggesting reduced turnover. BAS-1 synthesizes 5HT and DA from 5-hydroxytryptophan (5HTP) and L-DOPA (Fig. 2A), respectively, so this result suggests that these neurotransmitter intermediates are not sufficient to substitute in UPS regulation for their canonical biogenic amine products. The phenotype of these mutants, which also impair biogenic amine signaling, further support a role for these neurotransmitters in regulating UPS activity.

### Biogenic amine signaling promotes protein poly-ubiquitination without perturbing proteasome function

The stabilization of Ub^G76V^-GFP protein in biogenic amine signaling mutants could be due to either a reduction in its poly-ubiquitination or a reduction in its proteolysis by the proteasome. We made lysates from different biogenic amine synthesis mutants at L4+48h and then detected Ub^G76V^-GFP protein by Western blot either with anti-ubiquitin antibodies or anti-GFP antibodies (Fig. 3A). Little Ub^G76V^-GFP protein is detected in wild-type animals or *tbh-1* mutants at L4+48 h. Our control *skn-1(zj15)* mutants accumulated non-ubiquitinated, mono-ubiquitinated, di-ubiquitinated, and tri-ubiquitinated Ub^G76V^-GFP species, reflecting a reduction in polyubiquitination and/or the ability to clear polyubiquitinated substrates, consistent with reduced proteasome activity in these mutants (Fig. 3A). We observed a similar pattern of ubiquitinated Ub^G76V^-GFP accumulation in *cat-2*, *tdc-1*, and *tph-1* mutants. While we did not detect proteins larger than 70 kDa in biogenic signaling mutants or *skn-1(zj15)* using the anti-GFP antibodies, we did detect additional proteins ranging from 70 to 250 kDa in these mutants using anti-ubiquitin antibodies, which would detect endogenous ubiquitinated proteins in addition to the Ub^G76V^-GFP reporter. Although we cannot determine whether such proteins are mono-, di-, or poly-ubiquitinated, our results suggest that ubiquitinated forms of some endogenous proteins accumulate in these mutants. Together, these results suggest that DA, 5HT, and TA signaling promote either protein ubiquitination, proteolysis by the proteasome, or both.

**Figure 3.**
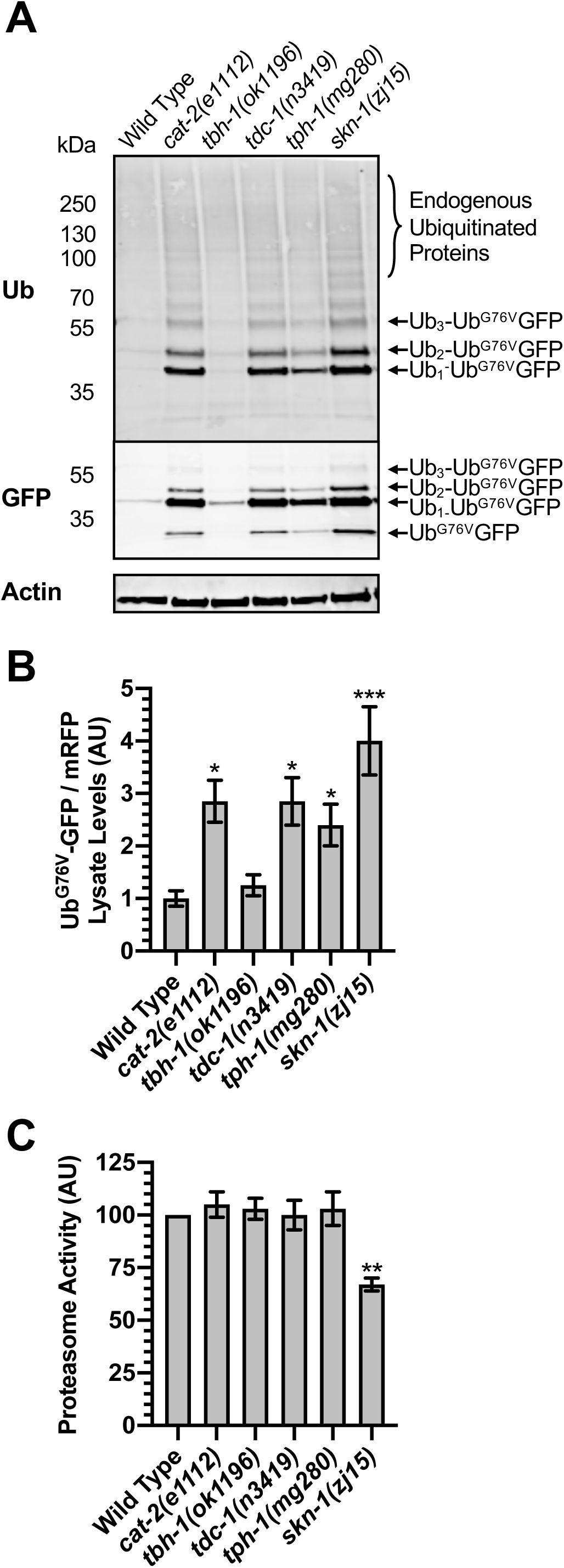
Biogenic amine signaling promotes protein poly-ubiquitination without perturbing proteasome function. A Western blot of lysed transgenic nematodes that express the Ub^G76V^-GFP and mRFP reporters in the hypodermis at the L4+48h stage, probed with antibodies recognizing ubiquitin, GFP, or actin as a loading control. The position of molecular weight markers is shown on the left of each blot. The position of Ub^G76V^-GFP protein, as well as Ub^G76V^-GFP with the indicated number of additional ubiquitin moieties, based on molecular weight, is indicated to the right of each blot. Twenty animals were loaded per lane for each indicated genotype. B Quantified fluorescence of Ub^G76V^-GFP normalized to mRFP from the lysates. *P<0.01, ***P<0.0001, ANOVA with Dunnetts posthoc comparison to wild type. N=3 trials. Error bars indicate SEM. C Quantified epoxomicin-sensitive proteasome activity (as measured through the turnover of fluorescent chymotrypsin substrate) from the same lysates in (B). **P<0.001, ANOVA with Dunnetts posthoc comparison to the empty RNAi vector control. N=3 trials. Error bars indicate SEM.

We also directly examined proteasome activity in biogenic signaling mutants. We generated lysates of L4+48h nematodes and first analyzed Ub^G76V^-GFP and mRFP levels in these lysates by fluorescence spectroscopy (Fig. 3B). We observed low levels of Ub^G76V^-GFP fluorescence at L4+48h in wild-type and *tbh-1* mutant animals, similar to what we observed by epifluorescence microscopy and Western blot analysis. By contrast, Ub^G76V^-GFP fluorescence was elevated in *cat-2*, *tph-1*, *tdc-1*, and *skn-1* mutants (Fig. 3B). We incubated the same lysates with the fluorescent chymotryptic substrate Suc-Leu-Leu-Val-Tyr-AMC to measure epoxomicin-sensitive proteasome activity. As expected, *skn-1(zj15)* control lysates showed reduced activity (Fig. 3C), which is in consonance with our quantification by epifluorescence microscopy (Fig. 1). By contrast, we did not observe a substantial change in proteasome activity in any of the biogenic amine mutants compared to wild type. When taken together, the accumulation of Ub^G76V^-GFP species with reduced ubiquitin valence (i.e., the shift from polyubiquitinated to non-ubiquitinated, mono-ubiquitinated, and di-ubiquitinated forms), combined with no observed change in proteasome activity in *cat-2*, *tph-1*, and *tdc-1* mutants, strongly suggests that DA, 5HT, and TA signaling does not regulate proteasome activity but instead either promotes the polyubiquitination of unstable proteins like Ub^G76V^-GFP or facilitates the ability of the proteasome to access such proteins.

### Biogenic amine signaling promotes expression of cytochrome P450 enzymes and eicosanoids

Biogenic amine signaling results in both transcriptional and post-translational changes to other signaling pathway components (Frederick & Stanwood, 2009, Klein, Battagello et al., 2019, Mohammad-Zadeh, Moses et al., 2008, Roeder, 2005). We reasoned that the long-term changes in UPS activity and proteostasis in biogenic amine mutants could reflect differences in gene expression. We compared the mRNA expression profile of the four biogenic amine mutants to wild-type animals during adulthood (L4+48 hours) using RNA-seq. We isolated poly-A(+) RNA from three replicates of each genotype, subjected each sample to whole genome sequencing (RNA-seq), aligned resulting reads to the *C. elegans* genome using STAR (Dobin, Davis et al., 2013), and assessed differential gene expression between genotypes using EdgeR and an FDR-adjusted q-value (Anders & Huber, 2010), focusing only on genes with significant changes (q<0.01, 2-fold or more, lower or higher, compared to wild type). Mutants for *cat-2* and *tdc-1* had the greatest number of differentially expressed genes (Fig. S1A,C), whereas *tph-1* and *tbh-1* mutants had only modest numbers of differentially expressed genes (Fig. S1B,D). Less than 1% of differentially expressed genes were commonly regulated in all four genotypes, and about 20% were commonly regulated in at least three genotypes (Fig. S1E,F). We used DAVID Functional Annotation Clustering enrichment analysis (Huang da, Sherman et al., 2009a, Huang da, Sherman et al., 2009b) to examine gene ontology and pathway enrichment for genes enriched in one or more of the genotypes (Fig. 4A). Although there was little gene ontology or pathway enrichment in genes that were overexpressed in any of the mutants relative to wild type, we observed enrichment of multiple ontology groups in genes that were underexpressed in one or more mutants. For example, genes involved in cuticle structure were enriched in downregulated genes in *tdc-1* mutants, suggesting a role for TA in cuticle formation and function. Multiple chloride channels were enriched in the downregulated genes found in both *cat-2* and *tdc-1* mutants, suggesting a role for DA and TA in modulating ion channel synthesis.

**Figure 4.**
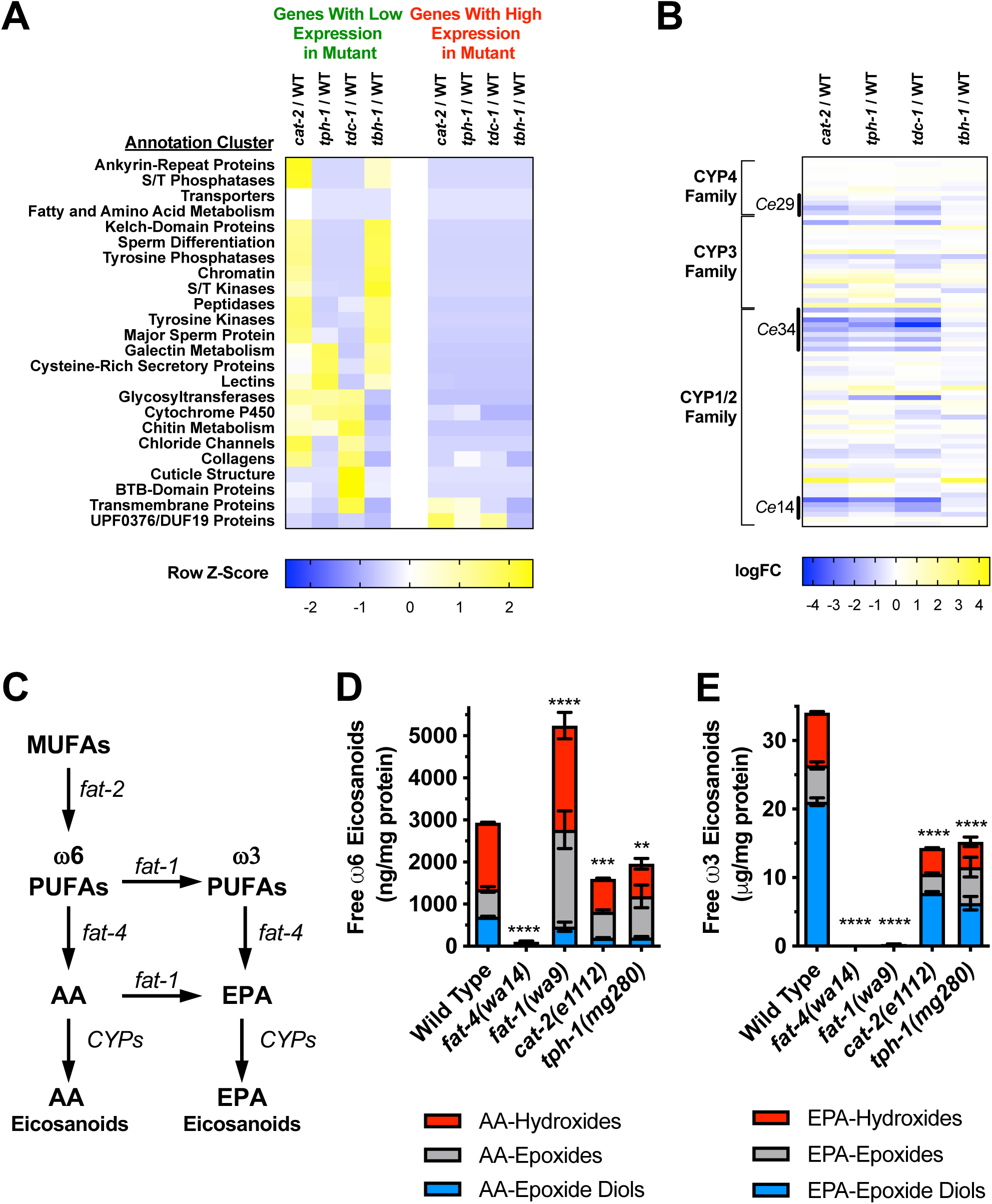
Biogenic amine signaling promotes expression of cytochrome P450 enzymes and eicosanoids. A Heat map of DAVID analysis showing enriched annotation clusters (e.g., gene ontology and pathways) in the genes showing differential expression between the indicated mutant and wild type. DAVID analysis of genes whose expression is lower in a mutant relative to wild type are highlighted with green text (four left hand columns), whereas DAVID analysis of genes whose expression is higher in a mutant relative to wild type are highlighted with red text (four right hand columns). Rows are labeled with each annotation cluster, with Z-scores (color scale at graph bottom) representing significant deviation between genotypes across the row for each annotation cluster. Annotation clusters (rows) are further clustered along the Y axis according to the Pearson average linkage method. B Heat map of log fold change (logFC, color scale at graph bottom) in the expression of *C. elegans* CYP genes between the indicated mutant and wild type. Genes whose expression is lower in a mutant relative to wild type are colored blue, whereas genes whose expression is higher in a mutant relative to wild type are colored yellow. Rows are labeled for the different CYP families and subfamilies. Specific genes (rows) are clustered according to sequence alignment (CLUSTAL W) to maintain the relationship of their molecular evolution. C Schematic representation of the synthesis pathway for mono-unsaturated fatty acids (MUFAs), poly-unsaturated fatty acids (PUFAs), arachidonic acid (AA), eicosapentaenoic acid (EPA), and their derived eicosanoids in *C. elegans*. The gene encoding each biosynthesis enzyme in the pathway is indicated next to the arrow representing its catalyzed reaction. D Abundance of the indicated free ω6 PUFA-derived eicosanoid, normalized to total protein, for each indicated genotype. Red, gray, and blue stacked bars indicate hydroxides, epoxide, and epoxide diol forms of eicosanoids, respectively. ****P<0.0001, ***P<0.001, **P<0.01, ANOVA with Dunnetts posthoc comparison to wild type. N=3 trials. Error bars indicate SEM. E Abundance of the indicated free ω3 PUFA-derived eicosanoid, as in (D).

As Ub^G76V^-GFP turnover was reduced in *cat-2*, *tph-1*, and *tdc-1* mutants, we were particularly interested in genes enriched in differentially expressed profiles from all three of these mutants. We noticed a large set of commonly-regulated genes between these three genotypes (21% of *cat-2* genes, 35% of *tph-1* genes, and 28% of *tdc-1* genes; Fig. S1E,F). These common genes were enriched for collagen and cuticle factors, chitin metabolism, glycosyltransferases, and cytochrome P450 monooxygenases (Fig. 4A), suggesting that one or more of these biological processes might be involved in a mechanism of proteostasis and UPS activity regulation shared by the three mutants.

We focused on cytochrome P450 proteins, which comprise a diverse superfamily of heme-containing enzymes involved in xenobiotic detoxification, fatty acid and steroid oxidation, and hormone synthesis and breakdown (Denisov, Makris et al., 2005, McLean, Sabri et al., 2005). The *C. elegans* genome contains 75 cytochrome P450 (CYP) genes assigned to 16 independent families within the CYP superfamily (Gotoh, 1998, Kulas, Schmidt et al., 2008). Most of the nematode CYP genes are similar to members of the mammalian CYP1/2, CYP3, and CYP4 families. We examined the expression of all *C. elegans* CYP genes in the mutants and found that the levels of the CYP14, CYP29, and CYP34 family members were underexpressed in *cat-2*, *tph-1*, and *tdc-1* mutants (Fig. 4B). These enzymes are closely related to the CYP2 and CYP4 microsomal monooxygenases, which produce eicosanoid signaling molecules and second messengers from long chain PUFAs (Capdevila, Falck et al., 2000, Kulas et al., 2008, McGiff & Quilley, 1999, Roman, 2002, Spector & Kim, 2015, Zeldin, 2001).

If DA, 5HT, and TA synthesis are required to promote the expression of CYP14, CYP29, and CYP34 genes, then do these biogenic amines promote the production of eicosanoids from PUFAs? *C. elegans* contains a complete set of genes to produce n-6 (ω6) and n-3 (ω3) long-chain PUFAs (Fig. 4C), including arachidonic acid (AA) and eicosapentaenoic acid (EPA), from oleic acid (OA)(Vrablik & Watts, 2013, Wallis, Watts et al., 2002, Watts & Browse, 2002, Ying & Zhu, 2016). We tested this hypothesis by collecting and lysing synchronized day 2 adult nematodes, followed by alkaline hydrolysis and LC-MS. AA-derived n-6 eicosanoids (Fig. 4D) were about 100 fold less abundant compared to EPA-derived n-3 eicosanoids (Fig. 4E), as previously observed (Kulas et al., 2008). As controls, we measured eicosanoid levels in *fat-1* mutants, which fail to synthesize n-3 long chain PUFAs, and *fat-4* mutants, which fail to synthesize AA and EPA from dihommogamma-linolenic acid (DGLA) and eicosatetraenoic acid (ETA) PUFAs, respectively (Watts & Browse, 2006). Mutants for *fat-4* lacked both AA- and EPA-derived eicosanoids (Fig. 4D,E). Mutants for *fat-1* lacked EPA-derived eicosanoids, but accumulated an unusually high level of AA-derived eicosanoids, perhaps due to the redirection of unconverted n-6 PUFAs into AA-derived eicosanoids. We focused on the best studied two biogenic amines, DA and 5HT, measuring eicosanoid levels in *cat-2* and *tph-1* mutants, respectively. We found a significant reduction in both AA- and EPA-derived eicosanoids in both mutants (Fig. 4D,E), indicating that DA and 5HT biogenic amines promote the synthesis of these lipid signaling molecules.

### The cytochrome P450 CYP-34A4 promotes UPS activity

Several mutations predicted to change conserved residues for one of these genes, *cyp-34A4* (Fig. 5A), were available from the Million Mutation collection (Thompson, Edgley et al., 2013), providing potential tools to analyze *cyp-34A4* function. The mRNA levels for *cyp-34A4* were significantly reduced in *cat-2*, *tph-1*, and *tdc-1* mutants, but not in *tbh-1* mutants (Fig. 5B). We mapped the amino acid changes for these mutations on an alignment between CYP-34A4 and the closely related rabbit CYP2C5 and human CYP2C8 (Fig. S2), for which there is extensive biochemical and structural data (Schoch, Yano et al., 2004, Williams, Cosme et al., 2000). The *gk933903* mutation changes a conserved proline to serine (P26S) in a loop that forms the membrane binding surface (Fig. 5A; Fig. S2). The *gk690081* mutation changes a conserved glycine to arginine (G395R) in the beta sheet structure that forms the membrane binding surface and helps coordinate lipid substrate interaction. The *gk759290* mutation changes a key conserved glycine to glutamate (G482E) at the end of a loop that helps stabilize helices that interact with lipid substrates. We introduced the *odIs77* transgene into these mutants and observed Ub^G76V^-GFP stabilization (Fig. 5C), similar to *cat-2*, *tph-1*, and *tbh-1* mutants, demonstrating that at least one of the CYP genes whose expression is elevated by biogenic amine signaling, is required to modulate UPS activity. As an independent confirmation, we used CRISPR/Cas9-mediated genome editing to introduce a small genomic rearrangement into *cyp-34A4*. This allele, *od107*, deletes 129 bases of the first two exons and the first intron, inserting 51 bases of novel DNA in its place. The rearrangement results in the deletion of amino acids 18-46, which encodes the N-terminal coil that forms the predicted membrane binding surface (Fig. 5A; Fig. S2). Following amino acid 17, *od107* creates two novel proline residues followed by a nonsense codon terminating translation of *cyp-34A4*, making this allele a likely null. We introduced *odIs77* into this mutant and observed stabilized Ub^G76V^-GFP in day 2 adults (Fig. 5C), similar to the other alleles of this gene. Taken together, our results demonstrate that DA, 5HT, and TA normally promote CYP-34A4 expression, and that CYP-34A4 is required for UPS activation in day 2 adults.

**Figure 5.**
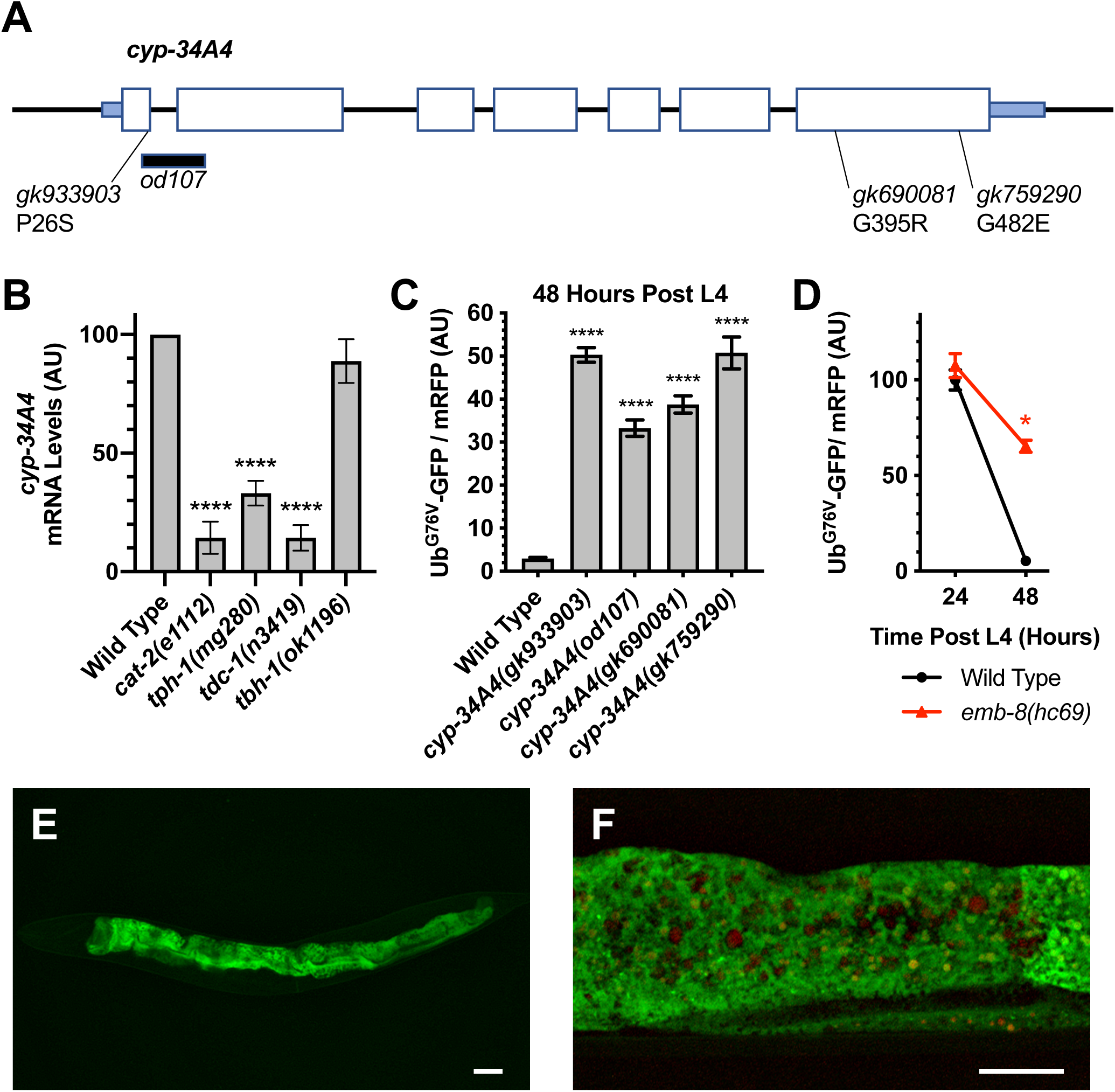
The cytochrome P450 CYP-34A4 promotes UPS activity. A Schematic of the *cyp-34A4* gene. Large boxes indicate coding sequences. Small boxes indicate UTRs. The location and nature of individual mutations is shown. Sequence deleted in *od107* is shown with a horizontal black bar. B Relative abundance of mRNA for the *cyp-34A4* gene. Values are based on qRT-PCR analysis, with the relative levels for each gene normalized to the value observed in the wild-type control. ***P<0.001, ANOVA with Dunnetts posthoc comparison to wild type. N=3 trials. Error bars indicate SEM. C Quantified fluorescence of Ub^G76V^-GFP normalized to mRFP in the hypodermis of animals of the indicated genotype at L4+48 hours. ****P<0.001, ANOVA with Dunnetts posthoc comparison to the wild-type control equivalent time point. N=20 animals per genotype and time point. Error bars indicate SEM. D. Quantified fluorescence of Ub^G76V^-GFP normalized to mRFP, as in (C). E Expression of the *cyp-34A4::gfp* transgene. Bar, 100 microns. F. Expression of the *cyp-34A4::gfp* transgene (green). Intestinal autofluorescence (red) can be distinguished. Bar, 100 microns.

Many cytochrome P450 enzymes interact with cytochrome P450 reductase, which allows electron transfer from NAD(P)H to reduce ferric P450 heme, to split oxygen, and then to hydroxylate substrate (McLean et al., 2005). EMB-8 encodes the lone *C. elegans* NADPH-cytochrome P450 reductase, and the mutation *emb-8(hc69)* is a loss of function mutation that gives a conditional embryonic lethal phenotype (Miwa, Schierenberg et al., 1980, Rappleye, Tagawa et al., 2003). We introduced the *odIs77* UPS reporter transgene into *emb-8* mutants, then hatched wild type and *emb-8* mutants with the transgene at a permissive temperature (15°C). When synchronized animals reached the L4 stage, we shifted to a restrictive temperature (20°C) and observed Ub^G76V^-GFP fluorescence. We found that Ub^G76V^-GFP was stable in day 2 adults (Fig. 5D), similar to *cyp-34A4* mutants. These results demonstrate that the cytochrome P450 reductase activity is required for UPS regulation, and that CYP-34A4 likely works with this reductase to oxidize a key substrate involved in that regulation.

To examine CYP-34A4 expression more closely, we generated a translational reporter containing the complete transcription unit and 2 kb of upstream sequence fused in frame to Venus at the CYP-34A4 C-terminus. We introduced this resulting transgene into the *C. elegans* germline and observed expression only in the intestine (Fig. 5E,F), with consistent expression through larval stages into adulthood, suggesting that CYP-34A4 might act non-autonomously to influence proteostasis in hypodermal tissues.

### PUFAs modulate UPS activity

If the enzymatic role of CYP-34A4 in UPS regulation is eicosanoid production rather than small molecule xenobiotic detoxification, then mutations that disrupt PUFA synthesis should cause the same Ub^G76V^-GFP stabilization phenotype as *cyp-34A4* and biogenic amine mutations. In *C. elegans*, PUFAs are generated from the mono-unsaturated fatty acid (MUFA) oleic acid (OA) by the Δ12-fatty acid desaturase FAT-2 (Fig. 6A), and *fat-2* mutants lack normal PUFAs (Peyou-Ndi, Watts et al., 2000, Watts & Browse, 2002). We introduced *odIs77* into *fat-2(ok873)* mutants, which have a deletion within *fat-2*, and observed stabilization of Ub^G76V^-GFP, similar to *cyp-34A4* mutants (Fig. 6B). In order to examine nematodes in which eicosanoids are selectively removed, we introduced *odIs77* into *fat-4(wa14)* mutants, which contain a nonsense mutation in the FAT-4 Δ5-fatty acid desaturase and lack AA, EPA, and all eicosanoids derived from these PUFAs (Fig. 4D,E; Fig. 6A) (Watts & Browse, 2002). We did not observe Ub^G76V^-GFP stabilization at L4+48h, instead observing precocious turnover of the UPS reporter at L4+24h (Fig. 4B). We also introduced *odIs77* into *fat-1(wa9)* mutants, which contain a nonsense mutation in the FAT-1 n-3-fatty acid desaturase, lack EPA and n-3 eicosanoids from EPA, and accumulate AA and n-6 eicosanoids (Fig. 4D,E; Fig. 6A) (Watts & Browse, 2002). We observed an even more dramatic precocious turnover of Ub^G76V^-GFP in these mutants (Fig. 6B). Double mutants between *fat-1* and *fat-4* resemble *fat-4* mutants (Fig. 6B), indicating that *fat-4* is epistatic to *fat-1*, and that the accumulation of AA or one of its eicosanoid derivatives might be responsible for the strong precocious turnover of Ub^G76V^-GFP in *fat-1* single mutants. Interestingly, *fat-3(ok1126)* mutants show a similar precocious turnover phenotype to that of *fat-4* mutants (Fig. 6B). These mutants contain a deletion for the FAT-3 Δ3-fatty acid desaturase, accumulate linoleic acid (LA) and alpha-linoleic acid (ALA), but fail to produce normal amounts of other PUFAs (Fig. 6A) (Watts & Browse, 2002). Taken together, our results suggest that (1) eicosanoids can play an instructive rather than a permissive role in UPS activity, as UPS activity can be driven either up or down depending on the eicosanoid profile, and (2) the full profile of PUFAs should be considered when assessing the role of lipid signaling in UPS activity.

**Figure 6.**
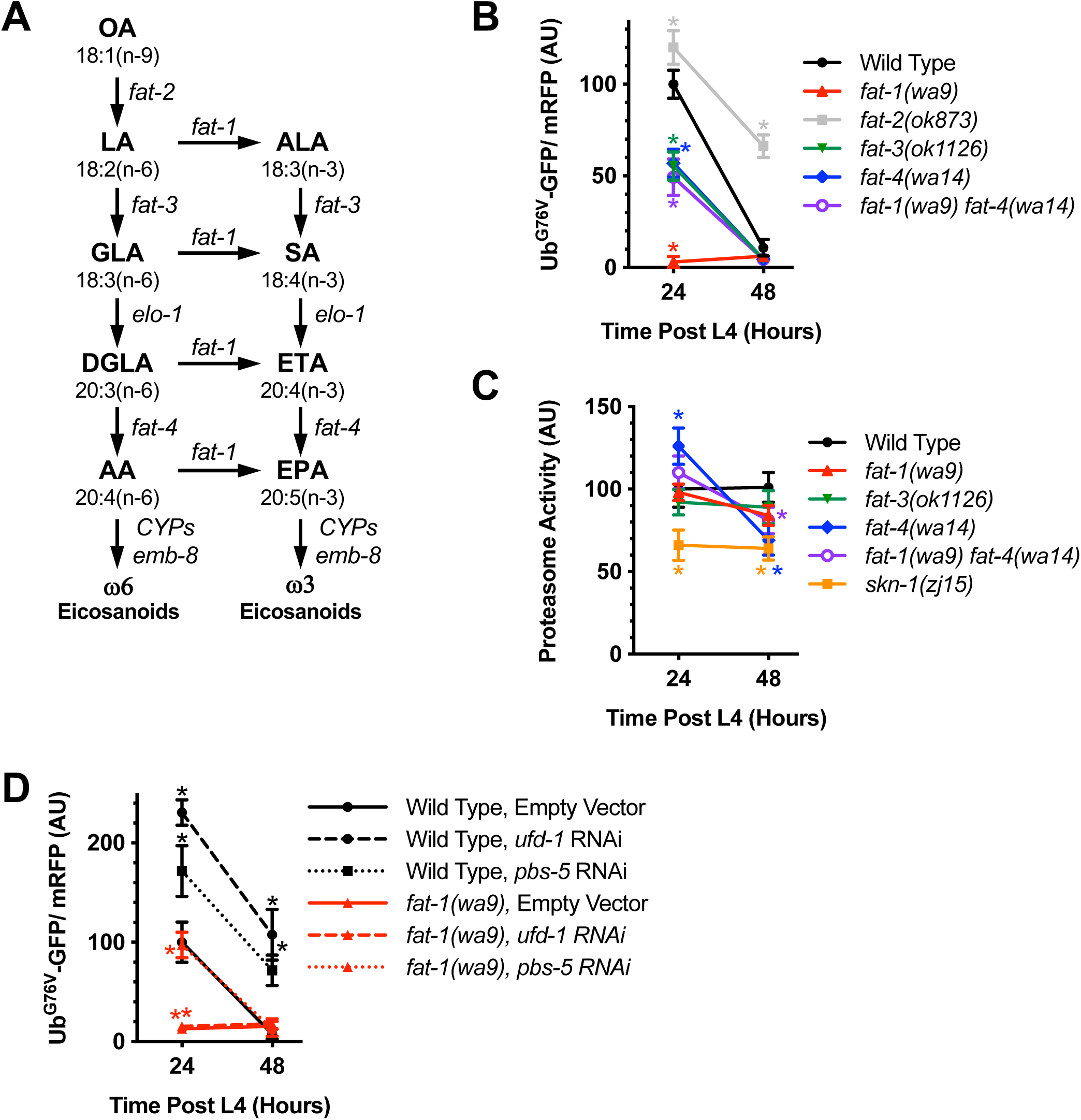
PUFAs modulate UPS activity. A Schematic representation of the PUFA and eicosanoid synthesis pathways in *C. elegans*. Carbon chain length, double bond number, and double bond location along the chain are indicated for each PUFA. The gene encoding each biosynthesis enzyme in the pathway is indicated next to the arrow representing its catalyzed reaction. B Quantified fluorescence of Ub^G76V^-GFP normalized to mRFP in the hypodermis of animals of the indicated genotype at the indicated time point. *P<0.001, ANOVA with Dunnetts posthoc comparison to the wild-type control equivalent time point. N=20 animals per genotype and time point. Error bars indicate SEM. C Quantified epoxomicin-sensitive proteasome activity (as measured through the turnover of fluorescent chymotrypsin substrate) from lysates of the indicated genotype and time point. *P<0.001, ANOVA with Dunnetts posthoc comparison to the wild-type control equivalent time point. N=3 trials. Error bars indicate SEM. D. Quantified fluorescence as in (B) for animals exposed to the indicated RNAi vector or empty vector control.

Is increased UPS activity responsible for the reduced levels of Ub^G76V^-GFP in *fat-1*, *fat-3*, and *fat-4* L4+24h mutants? Quantitative RT-PCR measurements of Ub^G76V^-GFP and mRFP mRNAs in these mutants showed no change relative to wild type, indicating that reduced Ub^G76V^-GFP levels are due to a post-transcriptional mechanism (data not shown). We generated lysates of L4+24h and L4+48h nematodes and incubated them with the fluorescent chymotryptic substrate Suc-Leu-Leu-Val-Tyr-AMC to measure epoxomicin-sensitive proteasome activity. As expected, *skn-1(zj15)* control lysates showed reduced activity (Fig. 6C). We did not observe a substantial change in proteasome activity in *fat-1* or *fat-3* mutants compared to wild type (Fig. 6C). However, *fat-4* mutants showed an elevated level of proteasome activity at L4+24h but a reduced level of activity at L4+48h (Fig. 6C), suggesting that one or more PUFAs or their eicosanoid derivatives can regulate proteasome activity.

If augmented UPS activity is responsible for precocious turnover of Ub^G76V^-GFP in mutants like *fat-1*, then a reduction in the UPS should suppress that turnover. Eggs harboring the *odIs77* transgene were hatched on bacterial lawns expressing RNAi constructs for either the UFD complex subunit *ufd-1*, the proteasome core particle subunit *pbs-5*, or an empty vector as control. As in Fig. 1, we found that Ub^G76V^-GFP levels in wild type were elevated relative to control in the *ufd-1* and *pbs-5* RNAi knockdown experiments by both day 1 and day 2 adulthood (Fig. 6D). Knockdown of *pbs-5* but not *ufd-1* resulted in some stabilization of Ub^G76V^-GFP in *fat-1* mutants at L4+24h, although the reporter in these mutants was soon degraded at L4+48h (Fig. 6D). These results suggest that overall UPS activity is increased in *fat-1* mutants, although as RNAi does not completely eliminate UFD complex or proteasome activity, we cannot rule out the activation of additional protein quality control mechanisms.

## DISCUSSION

Multicellular organisms use diverse and decisive pathways, including the UPS, to regulate proteostasis in response to extrinsic and intrinsic sources of proteotoxic stress; we propose that the *C. elegans* nervous system uses biogenic amine neurotransmitters and eicosanoid signaling to regulate proteostasis. Using a transgene expressing the metastable chimeric protein Ub^G76V^-GFP, which is a UPS substrate, we examined UPS activity in mutants that fail to synthesize different biogenic amines in *C. elegans*. We found that the biogenic amine neurotransmitters DA, 5HT, and TA promote the rapid Ub^G76V^-GFP turnover that occurs in epithelia as nematodes transition into fertile adulthood. We found that these biogenic amines are needed for the full expression of the CYP34 family of cytochrome P450 monooxygenases. CYP34 P450 enzymes produce eicosanoid signaling molecules from PUFAs, and we found that DA and 5HT are required for the full expression of eicosanoids. One of these biogenic amine-regulated P450s, *cyp-34A4*, is needed for the turnover of Ub^G76V^-GFP in adults, which matches the requirement for the biogenic amines that promote *cyp-34A4* expression. P450 monooxygenases produce eicosanoids with the assistance of cytochrome P450 reductases; as expected, Ub^G76V^-GFP turnover is impaired in mutants for the *C. elegans* reductase *emb-8*. Eicosanoids can be derived from n-3 or n-6 PUFAs, and we found that mutants impaired for all PUFA synthesis show impaired Ub^G76V^-GFP turnover. By contrast, mutants that synthesize n-6 but not n-3 PUFAs trigger the opposite phenotype, precocious Ub^G76V^-GFP turnover, suggesting that different eicosanoids have opposite roles in regulating proteostasis, and that eicosanoid signaling per se has an instructive rather than a permissive role in this regulation. Our results suggest that neurons use biogenic amine neurotransmitters as neurohormonal signals to modulate eicosanoid production from PUFAs, and that these eicosanoids in turn regulate proteostasis response pathways in non-neuronal tissues.

One of the key regulators of proteostasis in *C. elegans* is SKN-1, which has a dual function in embryonic development and in promoting the expression of xenobiotic detoxification enzymes in response to oxidative stress (An & Blackwell, 2003). Expression of proteasome subunit genes is reduced in *skn-1* mutants (Keith et al., 2016, Li et al., 2011, Niu et al., 2011). We therefore examined both Ub^G76V^-GFP turnover and measured proteasome activity in *skn-1(zj15)*, a fertile hypomorphic allele, and *skn-1(tm3411)*, a null deletion allele that results in sterile animals (Ruf et al., 2013, Tang et al., 2015). Ub^G76V^-GFP remained stable and proteasome activity was decreased in these mutants, consistent with decreased proteasome subunit expression. SKN-1 has traditionally been considered an ortholog of human Nrf2 because of their shared role in oxidative stress response. Although Nrf2 does not appear to regulate proteasome subunit expression, closely related Nrf1 does induce proteasome expression (Radhakrishnan 2010; Sha and Goldberg 2014;Steffen 2010). Thus, SKN-1 likely represents an ancestral form of the transcription factor, mediating both functions, and then diverged into two factors during mammalian evolution to handle each function separately.

Ub^G76V^-GFP levels change dramatically as animals transition into day 2 fertile adults, and multiple lines of evidence indicate that this reflects changes in UPS activity (Liu et al., 2011). The internal control mRFP is expressed from the same promoter and shares the same 5’UTR and 3’UTR regions as Ub^G76V^-GFP. Both reporters are expressed from the same integrated transgene. The levels of Ub^G76V^-GFP and mRFP mRNA do not change as animals enter fertile adulthood (Joshi et al., 2016). Mutating either key lysines on the Ub^G76V^-GFP transgene or knocking down proteasome or UFD complex activity prevents the decrease in Ub^G76V^-GFP in day 2 adults. Thus, it seems unlikely that the decrease in Ub^G76V^-GFP levels is due to decreases in reporter transcription or translation, but to changes in the ability of the UPS to recognize and degrade the reporter.

If the accumulation of Ub^G76V^-GFP in biogenic amine signaling mutants were due to a decrease in proteasome activity, then we should have detected a healthy decrease in our direct measurement of proteasome activity. Instead, the observed decrease in poly-ubiquitinated proteins and the unchanged activity in our proteasome assay strongly suggest that biogenic amine signaling does not regulate proteasome activity per se. One possibility is that biogenic amine signaling directly promotes the activity of the UFD complex rather than the proteasome. The physiological function of the UFD complex remains unclear, but it appears to act as an E4 ubiquitin ligase, recognizing proteins that have been mono-and poly-ubiquitinated by E3 ubiquitin ligases and amplifying their ubiquitin tagging through the addition of more ubiquitin moieties and the extension of the ubiquitin chain (Hoppe, 2005). Thus, the UFD complex provides an additional layer of global proteostasis control, and its regulation could determine the sensitivity of a cell to general proteotoxic stressors like oxidized or unfolded proteins. Consistent with this idea, mutants for UFD complex genes show reduced lifespan and sensitivity to proteotoxic stress (Liu et al., 2011, Mouysset, Kahler et al., 2006). We did not observe significant changes in the expression of UFD complex components in our analysis of biogenic amine mutants. However, it remains possible that biogenic amines promote UFD complex activity by a post-translational mechanism.

An alternative possibility is that biogenic amine signaling activates other proteostasis mechanisms, and that the stabilization of Ub^G76V^-GFP observed in the absence of biogenic amines is due to the UPS becoming overwhelmed handling the full proteostatic load left by the absence of these other mechanisms. In this model, proteasome activity pers se is not disrupted in the biogenic amine signaling mutants, consistent with our direct biochemical measurements of proteasome activity with fluorescent small molecule substrates. Rather the ability of the proteasome to obtain access to all ubiquitinated substrates *in vivo*, including our Ub^G76V^-GFP reporter, is disrupted. Our analysis of gene expression changes in biogenic amine mutants relative to wild type did not identify obvious alternative proteostasis pathways, but we did notice that biogenic amines were required to promote the expression of the CYP34 subfamily of P450s. The CYP34 subfamily is part of the larger CYP2 family, some members of which metabolize biogenic amine neurotransmitters, raising the possibility that the upregulation of CYP34 P450s by biogenic amines might act as a feedback, either positive or negative, to regulate biogenic amine signaling (Haduch & Daniel, 2018). Some mammalian CYP2 family members metabolize PUFAs into eicosanoids, and we observed a decrease in eicosanoid levels in *C. elegans* mutants defective for 5HT and DA synthesis. Thus, eicosanoids might be the link between these two biogenic amines and proteostasis.

Eicosanoids are produced from large chain PUFAs. Mutants that block the synthesis of all PUFAs resulted in the stabilization of Ub^G76V^-GFP, suggesting that one or more eicosanoid derivatives could regulate UPS activity and/or proteostasis. Such mutants are a bit extreme, as missing all PUFAs as well as their eicosanoid derivatives could result in observed phenotypes that are due to pleiotropic effects of losing so many of these molecules, which would cast PUFAs and eicosanoids in an indirect and permissive role in proteostasis. Surprisingly, *fat-1* mutants, which fail to produce n-3 PUFAs but synthesize augmented levels of n-6 PUFAs (and presumably their eicosanoid derivatives), show precocious Ub^G76V^-GFP turnover. This finding would suggest that the specific balance of eicosanoids can either increase or decrease UPS activity and/or proteostasis, which would cast PUFAs and eicosanoids in a more interesting instructive rather than permissive role in proteostasis.

The difference in phenotype when n-3 versus n-6 PUFA synthesis is disrupted would also suggest that different eicosanoids can have opposite effects on the same physiological process. Such disparate roles for different eicosanoids occur in well characterized functions for eicosanoids. For example, 20-HETE, an eicosanoid derived from n-6 PUFAs, has the opposite effect on blood pressure and heart arrhythmia observed for 17,18-EEQ, an eicosanoid derived from n-3 PUFAs (Fleming, 2011). The best understood “classic” eicosanoids are those derived from AA and EPA, yet *fat-4* mutants, which fail to make either of these PUFAs, still show some precocious Ub^G76V^-GFP turnover. However, eicosanoids are derived from other PUFAs besides just AA and EPA. For example, LA-derived eicosanoids include leukotoxin and iso-leukotoxin, which inhibit mitochondrial function and can trigger apoptosis in mammals (Sakai, Ishizaki et al., 1995, Stevens & Czuprynski, 1996). In *C. elegans*, eicosanoid derivatives of the PUFA DGLA induce germ cell death (Deline, Keller et al., 2015). Moreover, we cannot rule out the possibility that the true signaling molecules that regulate UPS activity are either novel molecules derived from classic eicosanoids through additional steps beyond P450, or nonclassical eicosanoids derived from PUFAs, similar to endocannabinoids (Cascio & Marini, 2015). Assigning specific physiological functions to specific eicosanoids will require a combined biochemical and genetic analysis of different PUFA synthesis enzyme and CYP mutants.

How might an eicosanoid signal regulate proteostasis? Eicosanoids bind to cell surface receptors like GPCRs and nuclear hormone receptors like PPAR (Brink, 2007, Marion-Letellier, Savoye et al., 2016). The *C. elegans* genome contains a diverse number of GPCRs and nuclear hormone receptors to match the diverse number of CYP genes, including the CYP families that produce eicosanoids. One or more of these genes might regulate proteostasis through transcriptional or post-translational means. Interestingly, n-6 PUFAs were shown in *C. elegans* to regulate lifespan through the activation of autophagy, another proteostasis mechanism that complements the UPS (O’Rourke, Kuballa et al., 2013). Different PUFA-derived signals might directly regulate the balance between autophagy and the UPS. Alternatively, changes in the PUFA profile might alter the total proteostatic load in a cell (e.g., by altering mitochondrial function and/or oxidative stress levels), either increasing or overwhelming UPS capacity and thereby increasing or decreasing protein turnover, respectively. Regardless of the specific mechanism, our findings raise the interesting possibility that the beneficial health effects of diets rich in fish oils, including n-3 and n-6 PUFAs, might be due in part to the regulation of proteostasis mechanisms like the UPS and autophagy.

## MATERIALS AND METHODS

### Strains and growth conditions

Standard methods were used to culture *C. elegans* (Brenner, 1974). Animals were grown at 20°C on standard NGM plates seeded with OP50 *Escherichia coli* unless otherwise indicated. The following strains were provided by the *Caenorhabditis* Genetics Center: *cat-2(e1112), cat-2(n4547), cyp-34A4(gk933903), cyp-34A4(gk690081), cyp-34A4(gk759290), tph-1(mg280), tdc-1(n3419), tdc-1(ok914), tbh-1(ok1196), tbh-1(n3243), fat-1(wa9), fat-2(ok873), fat-3(ok1126), fat-4(wa14), skn-1(zj15),* and *skn-1(tm3411).* Transgenic strains *odIs76[P_col−19_∷Ub^G76V^-GFP, Pcol−19∷mRFP]* and *odIs77[P_col−19_∷Ub^G76V^-GFP, Pcol−19∷mRFP]* have been described previously (Liu et al., 2011). When possible, researchers were blinded to the genotype of the observed sample.

### CRISPR/Cas9

CRISPR/Cas9 system was carried out using a CRISPR-Cas9-RNP injection mixture with the following final concentrations: *cyp-34A4* guide RNA (28 μM), Cas9 nuclease (2.5 μg/μl), *dpy-10* guide RNA (12μM), *dpy-10* repair oligo (0.5 μM), trans-activating crRNA (tracrRNA; 40 μM), KCl (25 mM), and HEPES, pH 7.4 (7.5 mM). These components were homogenously mixed by gentle pipetting and allowed to incubate at 37°C for 15 min. After incubation, the mixture was filtered and loaded into a microinjection pipette, and the gonads of 24 adults (adulthood day 1) were injected using the standard microinjection technique. Injected P0 worms were allowed to recover at 20°C. After three days, 51 Rol and/or Dpy worms were picked to individual plates, allowed to produce self-progeny. Genomic DNA of the F3 generation was extracted and screened by PCR for rearrangements at the *cyp-34A4 loci*. Candidate mutants were singled to NGM plates and confirmed by DNA sequencing of genomic PCR products. All knockout mutant animals were transferred to new plates and outcross for at least three generations to eliminate off targets.

### Statistics

Twenty animals were chosen per genotype so as to observe an effect size at least as large as the coefficient of variation. All data with normal distributions were analyzed with GraphPad Prism 8 in most cases using ANOVA with Dunnett’s post-hoc test or a Student t-test (two-tailed) with Holm-Sidek correction for multiple comparisons.

### RNA-seq analysis

Developmentally synchronized animals were obtained by hypochlorite treatment of gravid adults to release embryos. Synchronized embryos were hatched on NGM plates and grown at 20°C until 48h after the L4 stage of development. Fluorodeoxyuridine was used to prevent the development of second-generation embryos once animals reached fertile adulthood (Gandhi, Santelli et al., 1980). For each RNA-seq experimental replicate, populations were grown simultaneously under the same conditions. Total RNA from several large size NGM plates was isolated from animals using trizol (Invitrogen) combined with Bead Beater lysis in four biological replicates for each genotype. Yields ranged from 2.8 to 6.4 μg per sample. An mRNA library (single-end, 75-bp reads) was prepared for each sample/replicate using Illumina Truseq with PolyA selection (RUCDR). Libraries were sequenced on an Illumina HiSeq 2500 in Rapid Run Mode, resulting in high quality sequence reads (FAST QC Phred score average of 40 out to the full 75 bp length). Reads (30-40 million per genome and replicate) were mapped tp the C. elegans genome (WS245) and gene counts generated with STAR 2.5.1a. From 93-95% of reads uniquely mapped to the genome. Normalization and statistical analysis on gene counts were performed with EdgeR using generalized linear model functionality and tagwise dispersion estimates. An MDS plot revealed the fourth replicate for each genotype was subject to a strong batch effect, so the fourth replicate samples were dropped, and all analysis re-run using only replicates one, two and three from each genotype. Likelihood ratio tests were conducted for each biogenic amine mutant relative to wild-type *odIs77* in a pairwise fashion with a Benjamini and Hochberg correction. Lists of differentially expressed genes were analyzed by DAVID (v6.8, david.abcc.ncifcrf.gov/home.jsp) using functional annotation clustering and a low classification stringency as the method for assessing annotation enrichment. List comparisons and four set Venn diagrams were generated using Venny (Oliveros, 2007). RNA-seq data sets are available at NIH/NCBI GEO through accession number GSE145255.

### RNAi feeding

RNAi feeding protocols were as described previously (Timmons, Court et al., 2001)*. E. coli* (HT115) producing dsRNA for individual genes was seeded onto NGM plates containing 25 μg per ml carbenicillin and 0.2% lactose to induce the expression of the dsRNA for the gene of interest. The negative control was conducted by seeding the plates with HT115 containing empty vector pL4440. Synchronized embryos were hatched on onto each plate and grown at 20°C until 48h after the L4 stage of development. The respective clones contained sequences for *ufd-1* or *pbs-5* in RNAi vectors and were obtained from OpenBiosystems.

### Fluorescence microscopy, imaging analysis and intensity measurements

GFP- and mRFP-tagged fluorescent proteins were visualized in nematodes by mounting on 2% agarose pads with 10 mM tetramisole. Fluorescent images of *odIs76* and *odIs77* transgenic animals were observed using an AxioImager M1m (Carl Zeiss, Thornwood, NY). A 5 × (numerical aperture 0.15) PlanApo objective was used to detect GFP and mRFP signal. Imaging was done with an ORCA charge-coupled device camera (Hamamatsu, Bridgewater, NJ) by using iVision software (Biovision Technologies, Uwchlan, PA). Exposure times were chosen to capture at least 95% of the dynamic range of fluorescent intensity of all samples. GFP fluorescence was quantified by obtaining outlines of worms using images of the mRFP control. The mean fluorescence intensity within each outline was calculated (after subtracting away background coverslip fluorescence) for Ub^G76V^-GFP and mRFP signals using ImageJ. Fluorescent images of *cyp-34A4::gfp* transgenic animals were observed using a Chroma/89 North CrestOptics X-Light V2 spinning disk, a Chroma/89North Laser Diode Illuminator (405 nm and 470 nm lines to detect intestinal autofluorescence and GFP, respectively), and a Photometrics PRIME95BRM16C CMOS camera. Images were collected and analyzed with MetaMorph software.

### Measurement of 26S proteasome activity, GFP levels, and mRFP levels in lysates

Developmentally synchronized L4+24h and L4+48h staged animals were lysed in lysis buffer (50 mM HEPES, pH 7.5, 150 mM NaCl, 5 mM EDTA, 2 mM ATP, 1 mM dithiothreitol and protease inhibitor cocktail tablets; Roche) using Bead Beater. Lysates were transferred to microcentrifuge tubes and centrifuged at 15,000 rpm for 15 minutes at 4°C.

To measure proteasome activity, an equal amount of protein lysate (5 μg in 20 μl lysis buffer) was premixed with either 250 ng of proteasome inhibitor (epoxomicin, Boston Biochem, prepared in 50% dimethyl sulfoxide) or an equivalent volume of vehicle (50% dimethyl sulfoxide lacking the inhibitor), followed by the addition of proteasome assay buffer (200 μl; 25 mM HEPES, pH 7.5 and 0.5 mM ethylenediaminetetraacetic acid) containing a chymotryptic substrate (80 μM Suc-Leu-Leu-Val-Tyr -AMC; Boston Biochem). Reactions were incubated at 25°C for 1 h, and fluorescence (λex = 360 nm; λem = 465 nm) was detected after every 5 min using a Tecan Infinite F200 detector. Epoxomicin-sensitive activity was generated with triplicate measurements from two independent experiments, quantified, and plotted.

To measure GFP and mRFP in lysates, 5 μg of lysate in 100 μl of lysis buffer was placed in a 96-well plate, in duplicate. GFP and mRFP fluorescence was detected using a Tecan Infinite F200 detector using a filter set for either GFP (λex = 485 nm; λem = 535 nm) or mRFP (λexe = 590 nm; λem = 635). Relative fluorescence values were calculated.

### Western blotting and antibodies

Nematodes were synchronized by picking to fresh plates at L4 for *odIs77* and the various mutants. Protein samples (30 μl) were prepared from 20 synchronized nematodes for each genotype lysed at L4+48h. Equal amounts of protein samples were resolved by electrophoresis through 12% SDS polyacrylamide gel (Bio-Rad). Western blotting was performed using mouse anti-Ub (1:2000, Enzo), mouse anti-GFP (1:2000, Roche), and mouse anti-Actin (1:5000, Millipore). Western blotted proteins were visualized and quantified using fluorescent conjugated secondary antibodies from Odyssey and Licor Imaging system. Each experiment was repeated at least five times with lysates from separate nematode preparations.

### Endogenous eicosanoid measurements

The eicosanoid profile was determined for the wild-type as various mutant strains, with three independent cultures per strain. Nematodes collected at L4+48h were harvested as described for the RNA-seq experiments above, then prepared for LC–MS/MS analysis essentially as described previously (Kulas et al., 2008). Sample aliquots were mixed with internal standard compounds, including 19-HETE (hydroxyeicosatetraenoic acid), 20-HETE, 8,9-EET (epoxyeicosatrienoic acid), 11,12-EET, 14,15-EET, 5,6-DHET (dihydroxyeicosatrienoic acid), 8,9-DHET, 11,12-DHET, 14,15-DHET, 19-HEPE (hydroxyeicosapentaenoic acid), 20-HEPE, 8,9-EEQ (epoxyeicosatetraenoic acid), 11,12-EEQ, 14,15-EEQ, 17,18-EEQ, 5,6-DiHETE (dihydroxyeicosatetraenoic acid), 8,9-DiHETE, 11,12-DiHETE, 14,15-DiHETE, and 17,18-DiHETE (Cayman Chemicals), and then subjected to alkaline hydrolysis followed by solid-phase extraction of the metabolites. HPLC and MS conditions as well as the multiple reaction monitoring for the analysis of the CYP–eicosanoid profile were exactly as described previously (Arnold, Markovic et al., 2010).

## ACKNOWLEDGEMENTS

We thank Matthew Kiel and Iris Tu for their technical assistance in analyzing some of the ubiquitin reporter experiments. We thank A. Fire, the *C. elegans* Genetics Center, S. Mitani, and the Japanese National Bioresource Project for reagents and strains. We thank Brian Schubert and Daja O’Bryant for computational assistance. We thank Eunchan Park and Cathy Savage-Dunn for comments on the manuscript. The manuscript was funded by grants from the National Institutes of Health (R01 NS42023 and R01 GM101972, to CR), as well as the New Jersey Commission on Cancer Research and the Charles and Johanna Busch Fellowship (to KKJ); these agencies had no other role in the research or the manuscript.

## AUTHOR CONTRIBUTIONS

KKJ designed the genetic, molecular, behavioral, and cell biological experiments under the direct supervision of CR. CR, SP, and KKJ designed the RNA-seq analysis, and SP analyzed the results. CR, RM, and KKJ designed the eicosanoid biochemical analysis, and RM performed the mass spectrometry and analyzed the results. KKJ, TLM, and MK collected the data and analyzed the results from the remaining experiments under the direct supervision of CR. CR and KKJ wrote the manuscript.

## EXPANDED VIEW FIGURE LEGENDS

**Figure EV1**

A Volcano plot of differential gene expression between *cat-2* mutants and wild type. The log2 fold change (logFC) and false discovery rate are plotted along the X and Y axes, respectively. Genes that are significantly (P<0.01 FDR) underexpressed (>2 fold) in the mutant relative to wild type are indicated in green. Genes that are significantly overexpressed in the mutant are indicated in red.

B Volcano plot for *tph-1* relative to wild type, as per (A).

C Volcano plot for *tdc-1* relative to wild type, as per (A).

D Volcano plot for *tbh-1* relative to wild type, as per (A).

E Venn diagram of genes that are under-expressed in the indicated mutant relative to wild type. The number of genes in each category is indicated in each overlapping section of the diagram, along with the percentage the category comprises of the total list of differential genes for all four comparisons.

F Venn diagram of genes that are over-expressed in the indicated mutant, as per (E).

**Figure EV2**

CLUSTAL W alignment of *C. elegans cyp-34A4* and its closely related homologs 2C5 (in rabbit) and 2C8 (human). Structural elements based on known crystal structures are indicated by colored bars under the sequence. Substrate binding site residues (labeled with S) are in Helix A, Helix B, the B’-C loop, Helices F and F’, Helix G, Helix I, the loop between Helix K and Beta 2, and the loop following Helix G. Membrane binding surface residues (labeled with M) are found in Coil 1, Beta 1, and Beta 2. Residues coordinating heme (labeled with H) are in Helix C, Beta 2, and the loop before Helix G. Residues forming the interaction surface with NADPH Cytochrome P450 Reductase (labeled with R) are in Helix C, Helix G, and the loop before Helix G. The *gk933903* mutation changes a conserved proline in the loop that forms the membrane binding surface. The *gk690081* mutation changes a conserved glycine in the beta sheet structure that forms the membrane binding surface and helps coordinate lipid substrate interaction. The *gk759290* mutation changes a key conserved glycine at the end of a loop that helps stabilize helices F, F’, and G, which interact with lipid substrates. The *od107* mutation deletes parts of exons 1 and 2 and inserts 51 base pairs of novel sequence between the two. This results in the deletion of amino acids 18-46 (encoding the N-terminal coil that forms the membrane binding surface, as indicated by the horizontal line). Following amino acid 17, there are two novel proline residues followed by a nonsense codon.

